# N-acetyltransferase 2 genotypes amongst Zulu Speaking South Africans and isoniazid / *N*-acetyl-isoniazid pharmacokinetics during anti-tuberculosis treatment

**DOI:** 10.1101/864454

**Authors:** Thuli Mthiyane, James Millard, John Adamson, Yusentha Balakrishna, Cathy Connolly, Andrew Owen, Roxana Rustomjee, Keertan Dheda, Helen McIlleron, Alexander S. Pym

## Abstract

**Background:** Distribution of *N-acetyltransferase2* (*NAT2*) polymorphisms varies considerably among different ethnic groups. Information on *NAT2* single-nucleotide polymorphisms in South African population is limited. We investigated *NAT2* polymorphisms and their effect on isoniazid pharmacokinetics in Zulu black HIV-infected South Africans in Durban, South Africa. Methods: HIV-infected participants with culture-confirmed pulmonary tuberculosis (TB) were enrolled from two unrelated studies. Culture-confirmed participants were genotyped for *NAT2* polymorphisms 282C>T, 341T>C, 481C>T, 857G>A, 590G>A and 803A>G using Life Technologies pre-validated Taqman assays (Life Technologies, Paisley, UK). Participants underwent sampling for determination of plasma isoniazid and *N*-acetylisoniazid concentrations.

**Results:** Amongst the 120 patients, 63/120 (52.5%) were slow metabolisers (*NAT2*\*5/*5), 43/120 (35.8%) had intermediate (*NAT2*5/12*), and 12/120 (11.7%) had rapid genotype (*NAT2*4/*11, NAT2*11/12* and *NAT2*12/12*). NAT2 alleles in this study were *4, *5C, *5D, *5E, *5J, *5K, *5KA, *5T, *11A, *12A/12C and *12M. NAT2*5 was the most frequent allele (70.4%) followed by *NAT2*\*12 (27.9%). 34/40 had both PK results and *NAT2* genotyping results. The median area under the concentration-time-curve to infinity (AUC_0-∞_) interquartile range (IQR) was 7.81 (5.87 – 16.83) μg/ml/hr and maximum concentration (Cmax) 3.14 μg/ml (2.42 – 4.36) μg/mL. Individual polymorphisms were not equally distributed, with some represented in small numbers. Genotype did not correlate with phenotype, rapid genotype showing higher AUC_0-∞_ than slow but not significant, p=0.43.

**Conclusion:** There was high prevalence of slow followed by intermediate then rapid acetylator genotypes. The poor concordance between genotype and phenotype suggests that other factors or genetic loci influence INH metabolism, and warrants further investigation in this population.

## Introduction

Tuberculosis (TB) remains a leading cause of global morbidity and mortality, with approximately 10 million cases and 1.5 million deaths in 2018 (1). South Africa is a high TB burden country with an estimated 301,000 cases in 2018. The so-called ‘short-course’ treatment regimen recommended in international guidelines; consisting of 6 months of rifampicin and isoniazid, supplemented by pyrazinamide and ethambutol in the first 2 months, has remained largely unchanged for several decades. Whilst this regimen can achieve high relapse-free cure rates, a range of host and mycobacterial factors can influence treatment outcomes. There is increasing evidence that inter-individual variability in the pharmacokinetics (PK) of drugs within this regimen lead to heterogeneity in clinical outcomes(2, 3).

Pharmacogenomics describes one cause of PK variability due to polymorphisms in drug metabolising enzymes and transporters. During TB treatment, isoniazid is the paradigmatic case. Isoniazid is acetylated to its major metabolite, *N*-acetyl-isoniazid (AcINH), by the action of hepatic *N*-acetyltransferase 2 (*NAT2*). AcINH is subsequently rapidly hydrolysed to acetylhydrazine, which is also acetylated, to diacetyl-hydrazine, by the action of NAT2(4). Accumulated acetyl-hydrazine can be oxidised to form other, potentially hepatotoxic metabolites(4–6). Moreover, accumulated isoniazid can be metabolised by an alternative pathway where it is first hydrolysed to hydrazine, which has also been implicated in liver injury, before acetylation to acetyl-hydrazine, again by *NAT2*(4, 7). Hence, the activity of *NAT2* both dictates metabolism of isoniazid, and determines the availability of potentially hepatoxic hydrazine and acetyl-hydrazine metabolites. Within the 870-base pair *NAT2* gene, a number of low-activity single nucleotide polymorphisms (SNPs) have been characterised. The *NAT2* genotype has been shown to determine the rate of acetylation by *NAT2* in several populations(8). Individuals homozygous for the wild-type alleles are characterised as ‘rapid’ acetylators (RAs), those homozygous for low-activity SNPs as ‘slow’ acetylators (SAs) and heterozygotes as ‘intermediate’ acetylators (IAs)(9–13). SAs have a higher incidence of side-effects, particularly drug-induced hepatitis, during TB therapy, presumably due to higher levels of hepatoxic metabolites (14–20). Amongst the first-line TB drugs isoniazid has the greatest early bactericidal activity (EBA) and isoniazid PK parameters have been associated with rates of cure, sterilisation and acquired drug resistance(3, 21–27). A link between rapid acetylation and increased risk of poor treatment outcomes has been reported (28, 29).

*NAT2* genotype is known to differ amongst ethnic groups; with approximately 40-70% of Caucasians, Indians and African Americans characterised as SAs, versus only around 10% of Asian populations(30–42). *NAT2* genotype is not well characterised in the communities where TB is most prevalent, particularly in sub-Saharan Africa. South Africa has several black ethnic groups and few have been studied(43–45). Bach *et al* characterised 40% of a Zulu population as phenotypically slow acetylators but these findings have not been replicated, or informed by genotypic analysis(44). Moreover, South Africa has a high HIV prevalence and discordant relationships between *NAT2* genotype and isoniazid acetylator phenotype have been described amongst individuals living with HIV in other settings(46, 47).

We therefore characterised the relationship between *NAT2* genotype, isoniazid and AcINH PK and hepatotoxicity in a cohort of TB-HIV coinfected individuals in Durban, KwaZulu-Natal, South Africa.

## Methods

### Participants, study treatment and sample collection

Participants from two unrelated PK studies were included(48, 49). Both studies recruited black, Zulu-speaking adults living with HIV from KwaZulu-Natal, South Africa, between March 2007 and April 2010. Study 1 entitled “Bioavailability of the fixed dose formulation Rifafour containing isoniazid, rifampicin pyrazinamide, ethambutol and the WHO recommended first line anti-retroviral drugs zidovudine, lamivudine, efavirenz administered to new TB patients at different levels of immunosuppression.”. The results of this study have been previously reported (49). As shown in Table 2, for the purposes of this analysis, we used samples collected on day 1 of the study after an overnight fast, at pre-dose, 1, 2, 4, 5, 6, 8, and 12 hours post dose, with samples analysed for INH and AcINH for 60 participants with microbiologically proven pulmonary TB (positive sputum culture or smear) who received a standard first line TB regimen consisting of a FDC as described above. The INH dose was 150 mg, 225 mg, 300 mg and 375 mg per day for participants with weight 30-37 kg, 38 – 54 kg, 55 – 70 kg, and 70 kg and above, respectively, as per WHO guidelines(50). Each participant had blood collected on a paxgene tube for NAT2 genotyping. In addition, genotyping was performed on a further 20 participants without TB who were recruited to this study (49).

Study 2, entitled “Pharmacokinetics of Rifabutin Combined with Antiretroviral Therapy in the Treatment of Tuberculosis Patients with HIV Infection in South Africa”, was a randomised controlled trial of two different rifabutin doses co-administered with lopinavir/ritonavir-based antiretroviral therapy (51, 52). Participants initially received 6 weeks of standard intensive phase treatment, followed by 2 weeks with rifabutin 300mg daily replacing rifampicin. After 2 weeks of the continuation phase during which participants received only isoniazid and rifabutin (both 300mg daily) PK sampling was carried out. Individuals were fasted overnight, and a standard hospital breakfast served 2 hours after drug ingestion. Sampling was conducted pre-dose and at 2, 3, 4, 5, 6, 8, and 12 and 24 hours after drug intake, with samples analysed for isoniazid and AcINH for 40 participants. *NAT2* genotyping was performed on 40 participants with 34 participants having both PK sampling and genotyping.

All participants receiving anti-TB treatment in both studies were given pyridoxine for peripheral neuropathy prophylaxis and patients with CD4 counts below 200 cells/mm^3^ received cotrimoxazole. No participants were on antiretrovirals at the time of PK sampling. Both studies were approved by the South African Medicines Control Council (SAMCC), Biomedical Research Ethics Committee (BREC) of the University of KwaZulu-Natal (Study 1-E294/05; Study 2-BFC011/07) and the South African Medical Research Council (SAMRC) ethics committee. Study one was also approved by the WHO Ethics Research Ethics Committee. Written informed consent was obtained from all participants.

### NAT2 genotype procedures

Total Genomic DNA was isolated from whole blood using the QIAamp DNA mini kit (Qiagen, Crawly, UK) according to manufacturer’s instructions. Participants were genotyped, using the DNA Engine Chromo4 system (Bio-Rad Laboratories, Hercules, CA) and Opticon Monitor v.3.1 software (Bio-Rad Laboratories), for 6 *NAT2* SNPs; 282C>T, 341T>C, 481C>T, 857G>A, 590G>A and 803A>G using Life Technologies pre-validated probe-based Taqman assays as per manufactures instructions (Life Technologies, Paisley, UK). Each participant sample was analysed in duplicate.

### Haplotype assignment and acetylator genotype inference

Haplotype assignment from probe-based SNP data is poorly described in African populations. We elected to employ an unbiased PHASE analysis, which takes the dataset as a whole to assign the most likely haplotype for each individual, alongside a probability for this assignment (53, 54). Haplotype for each individual and acetylator genotype for each haplotype were defined as per the *NAT* gene nomenclature committee (55). Individuals with two rapid alleles were defined as RAs, those with two slow alleles as SAs and those with one fast and one slow allele as IAs.

### Isoniazid and *N-acetyl*-isoniazid PK and phenotype inference

Blood samples were collected and placed on ice immediately, before centrifugation within 60 minutes, immediate separation and storage of plasma at −70°C until analysis. Concentrations of isoniazid, AcINH and a 6-aminonicotinic acid internal control were quantified using validated high-performance liquid chromatography and tandem mass spectrometry (HPLC-MS/MS). Sample preparation included a protein precipitation with acetonitrile and subsequent dilution with water. Analytes were chromatographically separated using a Waters Exterra C18, 3.5μm, 50mm x 2.1mm column and detected using the AB Sciex 5500 Q-Trap mass spectrometer. All analytes were analysed isocratically with an acetonitrile/water/0.1% formic acid mobile phase. Isoniazid, AcINH and the internal standard were analysed at mass transitions of the precursor ions (m/z) 137.9, 180.1 and 138.7 to the product ions (m/z) 66.0, 78.6 and 50.9, respectively. Chromatographic data acquisition, peak integration and quantification of analytes was performed using Analyst^®^ software version 1.5.2. We constructed time-concentration curves in the *PK* package in *R* for windows (version 3.5.1). We characterised the isoniazid and AcINH PK parameters maximum concentration (C_max_), time to maximum concentration (T_max_), area under the concentration curve from zero to infinity (AUC_0-∞_), apparent oral clearance (CL) and elimination half-life and compared C_max_ to published efficacy targets (56). AUC_0-∞_ was calculated using the trapezoid rule, apparent oral clearance estimated by dose / AUC_0-∞_ and elimination half-life by regression analysis of log_10_ concentrations of the terminal exponent of elimination. We analysed the ratio of log_10_ AcINH to log_10_ isoniazid at two and four hours to assess acetylation phenotype.

Sample processing and HPLC-MS was initially conducted in 2010 for Study 1. Samples remained in storage and were later moved to a new storage facility before they were shipped to a different laboratory for determination of isoniazid and AcINH concentrations as above (having previously only had isoniazid concentrations determined). To confirm the Integrity of these samples we compared the isoniazid AUC_0-∞_ of the current analysis with that previously reported on the same samples analysed in 2010.

### Statistical methods

All data were entered in Epidata and transferred to either Stata (version 14) or *R* for windows (version 3.5.1) for statistical analysis. Demographic characteristics were presented as frequencies and percentages for categorical variables, and as means with standard deviations for continuous variables. Descriptive PK data was described as median and inter-quartile ranges. C_max_ and AUC_0-∞_ were log-transformed prior to comparison between genotypes. PK parameters were compared, by genotype, using the Wilcoxon rank-sum test or Kruskal–Wallis test.

### Hepatic adverse events

Hepatic adverse events were defined as elevated alanine transaminase (ALT) and aspartate transaminase (AST), elevated alkaline phosphatase and elevated total bilirubin, graded as per Division of AIDS toxicity table for grading severity of HIV-positive adult adverse events.

## Results

### Participant characteristics

One hundred and twenty-two individuals living with HIV participating in two PK studies were included in the study. Eighty participants in study 1 were included in the *NAT2* genotyping analysis and 60 in the PK analysis (with 58 individuals having both PK and genotype data), while 40 participants in study 2 were included in the PK analysis and 40 participants included in *NAT2* genotyping analysis (with 34 individuals having both PK and genotype data). Key characteristics are outlined in table 1. Participants in study 1 included 60 with pulmonary TB and HIV co-infection; 40 with CD4 count >200 cells/mm^3^ and, 20 with CD4 count <200 cells/mm^3^ as well as 20 participants living with HIV and without TB (who contributed only genotype data). All 40 participants in study 2 had TB and HIV coinfection, with a CD4 count of 200 cells/mm^3^ or below. In the combined studies, 66.7% of participants had CD4 counts <200 cells/mm^3^ and 33.3% had CD4 count >200 cells/mm^3^. The Median age was 33.1 years (IQR 18-53). Only 15 (12.5%) of patients had a BMI < 18.86 kg/m^2^.

**Table 1:** Demographic characteristics.

| **Table 1: Demographic characteristics** |  |  |  |
| --- | --- | --- | --- |
|  | **Study 1 (n=80)** | **Study 2 (n=40)** | **Overall (n=120)** |
| Demographics |  |  |  |
| Median age (range) | 33 (18-48) | 33.6 (24-53) | 33.1 (18-53) |
| Male sex (%) | 36 (45%) | 24 (60.0%) | 60 (50%) |
| Zulu ethnicity (%) | 80 (100%) | 40 (100%) | 120 (100%) |
| Mean weight (SD) | 58.7 (11.9) | 58.9 (9.7) | 58.7 (11.2) |
| Mean BMI (SD) | 23.0 (5.2) | 23.1 (3.9) | 23.1 (4.8) |
| BMI <18.5 (%) | 13 (16.3%) | 2 (5.0%) | 15 (12.5%) |
| Median CD4 (range) | 210.5 (10-500) | 128 (61 – 199) | 161 (10-500) |
| CD4 < 200 (%) | 40 (50%) | 40 (100%) | 80 (66.7%) |

### *NAT2* genotype and deduced phenotype

One hundred and twenty participants (80 from study 1 and 40 from study 2) were genotyped. Allele and haplotype frequencies and deduced phenotypes are outlined in tables 2–5. We identified 12 different alleles in the population. The most common allelic group was *NAT2*5* (70.4%) followed by *NAT2*12* (27.9%). From the *NAT2*5* group *NAT2*5C* (21.3%), *NAT2*5J* (17.5%), *NAT2*5D* (14.6%) and *NAT2*5K* (10.4%) were the most common. The *NAT2*12* group was predominantly *NAT2*12C*. The deduced phenotype was 11.7% rapid, 35.8% intermediate and 52.5% slow.

**Table 2.** PK time points and dosing. Pharmacokinetic time points and dosing

| Study | Schedule of pharmacokinetic sampling (day of TB treatment) | Treatment |  |  |  |
| --- | --- | --- | --- | --- | --- |
| Study 1 | Days 1; with sampling pre-dose and at 1, 2, 4, 6, 8, 12 hours after the dose. | 4 drug FDC formulation (EMB/ RMP/ INH/ PZA 275/150/75/400 mg) dosed daily by weight band: |  |  |  |
|  |  | Weight in kilograms |  |  |  |
|  |  | 30-37 | 38-54 | 55-70 | > 70 |
|  |  | 2 tablets | 3 tablets | 4 tablets | 5 tablets |
| Study 2 |  | Enrolment – week 6: standard weight band based treatment with RMP, INH, PZA and EMB (as in study 1) |  |  |  |
|  |  | Week 6 & 7: RMP replaced with RFB 300 mg daily |  |  |  |
|  | Day 63 (after 2 weeks on continuation phase RBN+INH) with sampling pre-dose and 2, 3, 4, 5, 6, 8, 12 and 24 hours after the dose. | week 8 & 9: RFB 300 mg/INH 300 mg |  |  |  |

PK= pharmacokinetics; RMP=rifampicin; PZA=pyrazinamide; EMB=ethambutol; FDC=fixed dose combination

**Table 3.** NAT2 diplotypes and genotypes and deduced phenotype in the study group.

| Observed<br>Diplotype† | n | Genotype | Phenotype |
| --- | --- | --- | --- |
| -20000 | 1 | 5D/5K | SLOW |
| 000020 | 1 | 12A/12A | RAPID |
| 001000 | 1 | 4/11A | RAPID |
| 001020 | 6 | 12A/12C | RAPID |
| 002010 | 2 | 11A/12C | RAPID |
| 002020 | 4 | 12C/12C | RAPID |
| 01-020 | 1 | 5C/12C | INTERMEDIATE |
| 010010 | 1 | 5D/12A | INTERMEDIATE |
| 010020 | 2 | 5C/12A | INTERMEDIATE |
| 010110 | 2 | 5E/12A | INTERMEDIATE |
| 011010 | 3 | 5D/12C | INTERMEDIATE |
| 011020 | 15 | 5C/12C | INTERMEDIATE |
| 011110 | 3 | 5E/12C | INTERMEDIATE |
| 0200-0 | 1 | 5C/5D | SLOW |
| 020000 | 1 | 5D/5D | SLOW |
| 020010 | 10 | 5C/5D | SLOW |
| 020020 | 3 | 5C/5C | SLOW |
| 020100 | 3 | 5D/5E | SLOW |
| 020110 | 1 | 5C/5E | SLOW |
| 110010 | 1 | 5K/12A | INTERMEDIATE |
| 110110 | 1 | 5K/12C | INTERMEDIATE |
| 111010 | 5 | 5K/12C | INTERMEDIATE |
| 111020 | 1 | 5T/12C | INTERMEDIATE |
| 111110 | 6 | 5J/12C | INTERMEDIATE |
| 120000 | 7 | 5D/5K | SLOW |
| 120010 | 5 | 5C/5K | SLOW |
| 120011 | 1 | 5C/5KA | SLOW |
| 120020 | 1 | 5C/5T | SLOW |
| 120100 | 7 | 5D/5J | SLOW |
| 120110 | 8 | 5C/5J | SLOW |
| 120200 | 1 | 5E/5J | SLOW |
| 211020 | 1 | 5T/12M | INTERMEDIATE |
| 211110 | 1 | 5J/12M | INTERMEDIATE |
| 220001 | 1 | 5K/5KA | SLOW |
| 220100 | 4 | 5J/5K | SLOW |
| 220110 | 1 | 5J/5T | SLOW |
| 2202-0 | 1 | 5J/5J | SLOW |
| 220200 | 6 | 5J/5J | SLOW |

†Observed diplotypes are shown as the number of mutations identified in each individual for each SNP. 0 = wild type, 1 = heterozygous, 2 = homozygous, - = blank. The SNP order is 282, 341, 481, 590, 803, 857.

**Table 4.** Frequency of NAT2 alleles in the study group.

| Table 4. Frequency of NAT2 alleles in the study group. |  |  |
| --- | --- | --- |
| Allele designation | n | % |
| NAT2*4 | 1 | 0.4 |
| NAT2*5 | 169 | 70.4 |
| NAT2*11 | 3 | 1.3 |
| NAT2*12 | 67 | 27.9 |
| Total | 240 | 100 |

**Table 5.** Frequency of NAT2 alleles.

| Table 5. Frequency of NAT2 alleles. |  |  |
| --- | --- | --- |
| Allele | n | % |
| NAT2*4 | 1 | 0.4 |
| NAT2*5C | 51 | 21.3 |
| NAT2*5D | 35 | 14.6 |
| NAT2*5E | 10 | 4.2 |
| NAT2*5J | 42 | 17.5 |
| NAT2*5K | 25 | 10.4 |
| NAT2*5KA | 2 | 0.8 |
| NAT2*5T | 4 | 1.7 |
| NAT2*11A | 3 | 1.3 |
| NAT2*12A | 14 | 5.8 |
| NAT2*12C | 51 | 21.2 |
| NAT2*12M | 2 | 0.8 |
| Total | 240 | 100 |

### Isoniazid and *N-acetyl*-isoniazid PK

As above, to assess sample integrity for Study 1 we compared the AUC_0-∞_ of the current analysis with that previously reported on the same samples analysed in 2010. The median (IQR) AUC_0-∞_ was 5.53 (3.63 – 9.12), processed at University of Cape Town (UCT) in 2009 and 5.70 (3.85 – 7.94), processed at Africa Health Research Institute (AHRI) laboratory in 2014, suggesting that the integrity of the samples was maintained for isoniazid, but cannot be confirmed for AcINH.

Study 1 showed rapid absorption, with a median (IQR) isoniazid T_max_ of 1 hr(1 - 2). Isoniazid exposure was variable amongst individuals with median (IQR) C_max_ 1.47 (1.14 – 1.85) μg/ml and AUC_0-∞_ 5.53 (3.63 – 9.12) μg.h/ml. Median (IQR) elimination half-life was relatively slow at 2.27 (1.69 – 3.56) h. We compared these isoniazid PK measures to published targets; 98.28% (57/58) failed to attain the minimum 2-hour plasma concentration target of 3 μg/ml (56). PK parameters by genotype are shown in table 8(A), unexpectedly median half-life was slowest, apparent oral clearance lowest and AUC_0-∞_ highest amongst genotypically rapid acetylators, with the reverse true for genotypically slow acetylators, although none of these differences was statistically significant. Similarly, there were no statistically significant differences by genotype for AcINH C_max_, elimination half-life or AUC_0-∞_. Median isoniazid and AcINH time-concentration curves are given in Figure 1(A).

**Figure 1A:**
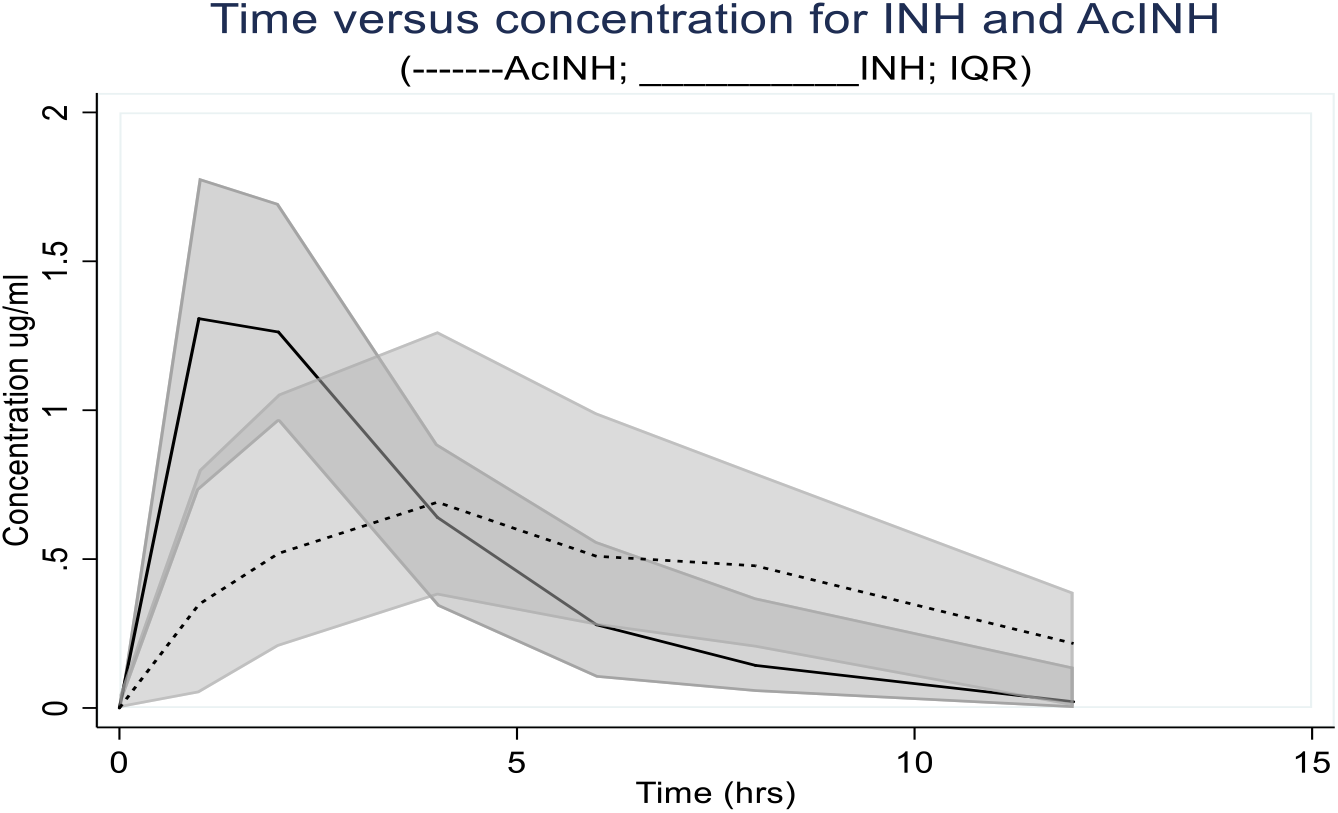
Study 1 median INH and AcINH concentration over time for INH and AcINH for 58 patients. Shaded area; IQR.

Absorption was rapid in Study 2, with a median INH T_max_ of 2 hrs. INH exposure was also variable amongst individuals with median (IQR) C_max_ 3.14 μg/ml (2.39 – 4.34) and AUC_0-∞_ 10.76 μg.hr/ml (8.24 – 28.96 μg/ml). Median elimination half-life was 2.62hr (2.26 – 4.07). Again, we compared these INH PK measures to published PK targets; 47.5% (19/40) failed to attain the minimum 2-hour plasma concentration target of 3 μg/ml. PK parameters by genotype are shown in table 8(B). For both isoniazid and AcINH and across the PK parameters; C_max_, AUC_0-∞_ and elimination half-life, variability (both range and IQR) were increased amongst those genotyped as SAs. Again however, there were no statistically significant differences between these PK parameters by genotype. Median isoniazid and AcINH time-concentration curves are given in Figure 1(B).

**Figure 1B:**
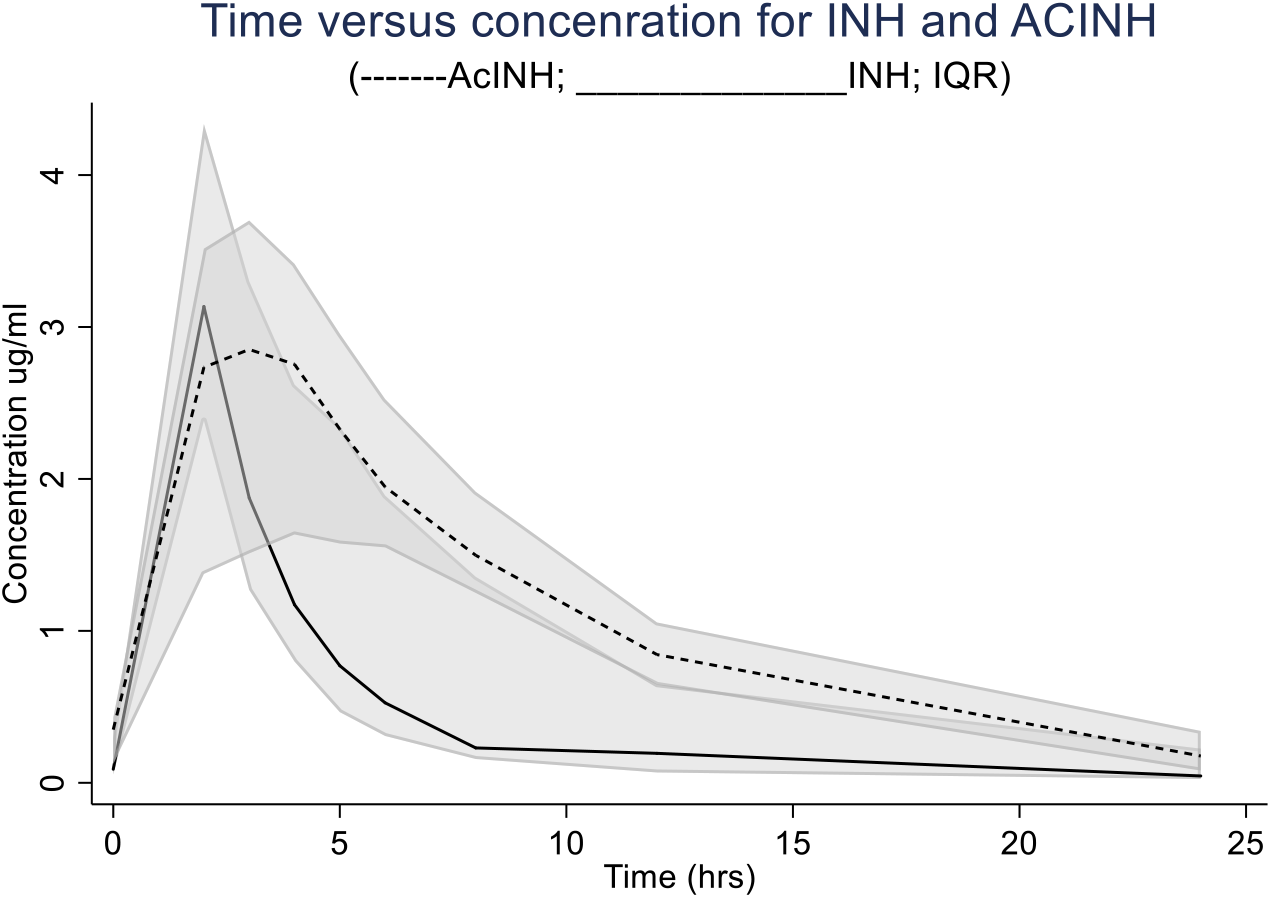
Study 2 median INH and AcINH concentration over time for INH and NA-INH for 34 patients. Shaded area: IQR.

**Table 6:** Frequency distribution of NAT2 genotypes and deduced phenotype in the study group.

|  |  |  |  |
| --- | --- | --- | --- |
| Genotype | n | % | Acetylator status |
| NAT2*4/*11 | 1 | 0.8 | RAPID |
| NAT2*12/*12 | 11 | 9.2 |  |
| Nat2*11/*12 | 2 | 1.7 |  |
| NAT2*5/*12 | 43 | 35.8 | INTERMEDIATE |
| NAT2*5/*5 | 63 | 52.5 | SLOW |
| Total | 120 | 100 |  |

**Table 7:**
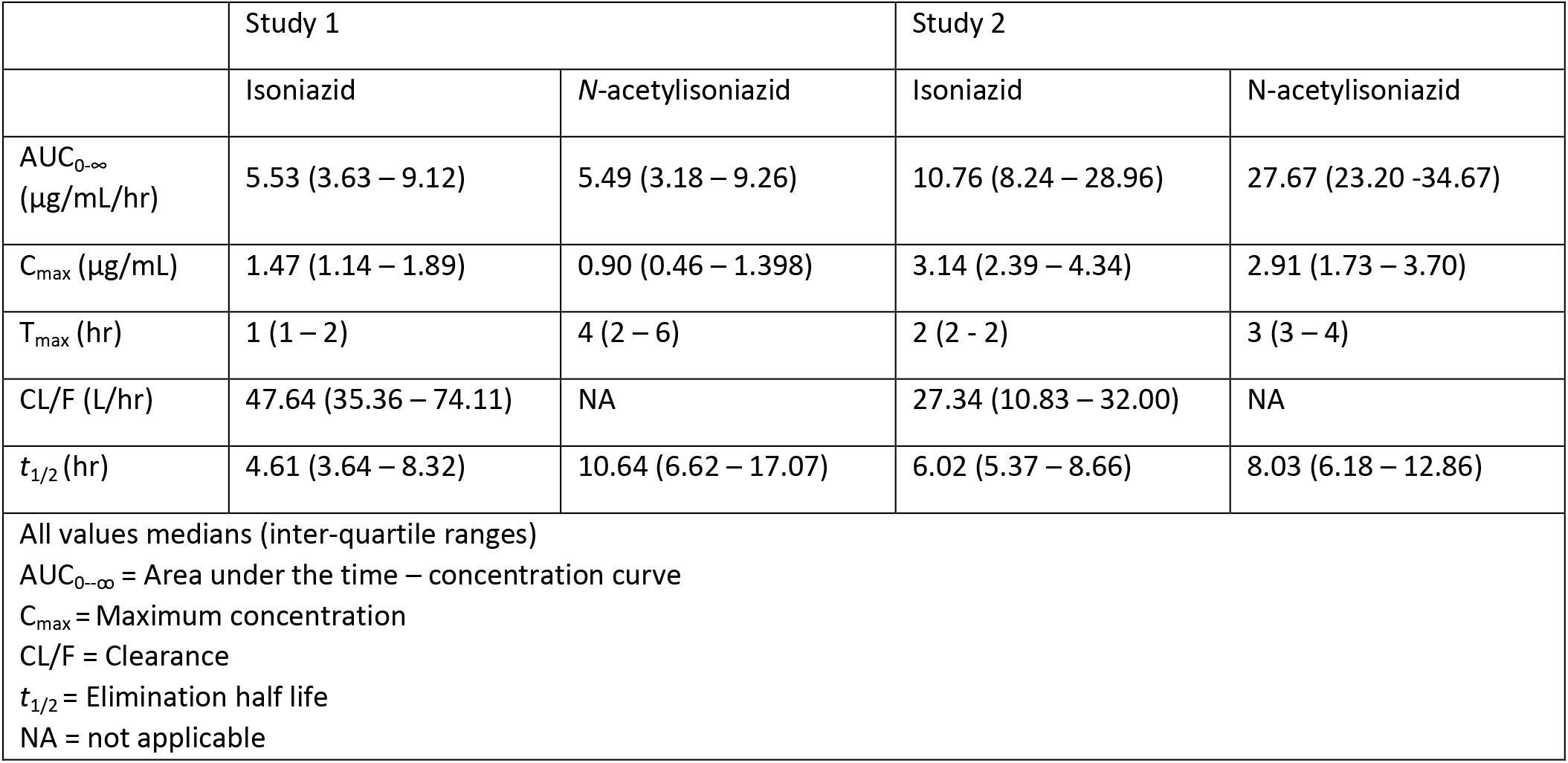
Overall isoniazid and N-acetyl-isoniazid PK.

**Table 8(A):** Study 1 PK parameters by genotype.

|  | Isoniazid |  |  | N-acetyl-Isoniazid |  |  |
| --- | --- | --- | --- | --- | --- | --- |
|  | Slow | Intermediate | Rapid | Slow | Intermediate | Rapid |
| AUC<sub>0-∞</sub> (µg/mL/hr) | 5.34 (3.44 – 7.93) | 6.04 (4.27 – 7.53) | 7.56 (5.99 -9.60) | 5.71 (4.19 – 11.01) | 7.34 (3.15 – 10.9) | 2.81 (0.55 – 5.05) |
| C<sub>max</sub> (µg/mL) | 1.47 (0.97 – 1.89) | 1.54 (1.25 – 1.76) | 1.42 (1.20 -2.05) | 0.94 (0.63 – 1.68) | 1.07 (0.49 – 1.70) | 0.38 (0.90 – 0.90) |
| T<sub>max</sub> (hr) | 1 (1 -2) | 1 (1 – 2) | 2 (2 – 2) | 4 (2 – 4) | 4 (4 – 7) | 6 (4 -6) |
| CL/F (L/hr) | 57.05 (37.84 – 103.56) | 43.53 (32.05 – 64.33) | 37.75 (31.27 – 47.92) | NA | NA | NA |
| t<sub>1/2</sub> (hr) | 4.67 (3.64 – 8.32) | 4.00 (3.35 – 5.19) | 8.56 (5.69 – 14.44) | 9.42 (5.75 – 17.07) | 6.55 (6.68 – 10.93) | 14.78 (10.65 – 22.41) |
| All values medians (inter-quartile ranges)<br>AUC<sub>0-∞</sub> = Area under the time – concentration curve<br>C<sub>max</sub> = Maximum concentration<br>CL/F = Clearance<br>t<sub>1/2</sub> = Elimination half life<br>NA = not applicable |  |  |  |  |  |  |

**Table 8(B):** Study 2 PK parameters by genotype.

|  | Isoniazid |  |  | N-acetyl-Isoniazid |  |  |
| --- | --- | --- | --- | --- | --- | --- |
|  | Slow | Intermediate | Rapid | Slow | Intermediate | Rapid |
| AUC<sub>0-∞</sub> (µg/mL/hr) | 10.76 (9.73 -31.21) | 9.09 (7.3 -18.75) | 26.99 | 26.04 (22.99 -32.76) | 6.28 (5.25 -10.01) | 28.53 |
| C<sub>max</sub> (µg/mL) | 3.47 (2.49 - 4.49) | 2.96 (2.33 - 4.02) | 3.94 | 2.85 (1.52 - 3.68) | 3.28 (2.53 - 4.01) | 1.91 |
| T<sub>max</sub> (hr) | 2 (2 – 2) | 2 (2 – 2) | 2 | 3 (3-4) | 3 (3 – 3) | 4 |
| CL/F (L/hr) | 27.87 (9.66 -30.83) | 33.33 (16.01 - 41.17) | 11.12 | NA | NA | NA |
| t<sub>1/2</sub> (hr) | 4.61 (3.9 -5.34) | 4.46 (3.9 - 7.88) | 8.28 | 5.81 (4.9 - 7.25) | 6.28 (5.25 -10.01) | 10.97 |

All values medians (inter-quartile ranges)
AUC<sub>0-∞</sub> = Area under the time – concentration curve
C<sub>max</sub> = Maximum concentration
CL/F = Clearance
t<sub>1/2</sub> = Elimination half life
NA = not applicable

For both studies we calculated the log_10_ AcINH: log_10_ isoniazid ratio, as a measure of acetylation, at two and four hours post-dose and analysed this ratio by genotype (figure 2 & 3). In both studies we saw no statistically significant difference in ratios between genotypes at either two or four hours. In Study 2 we again saw increased variability in this metric amongst those genotyped as SAs.

**Figure 2:**
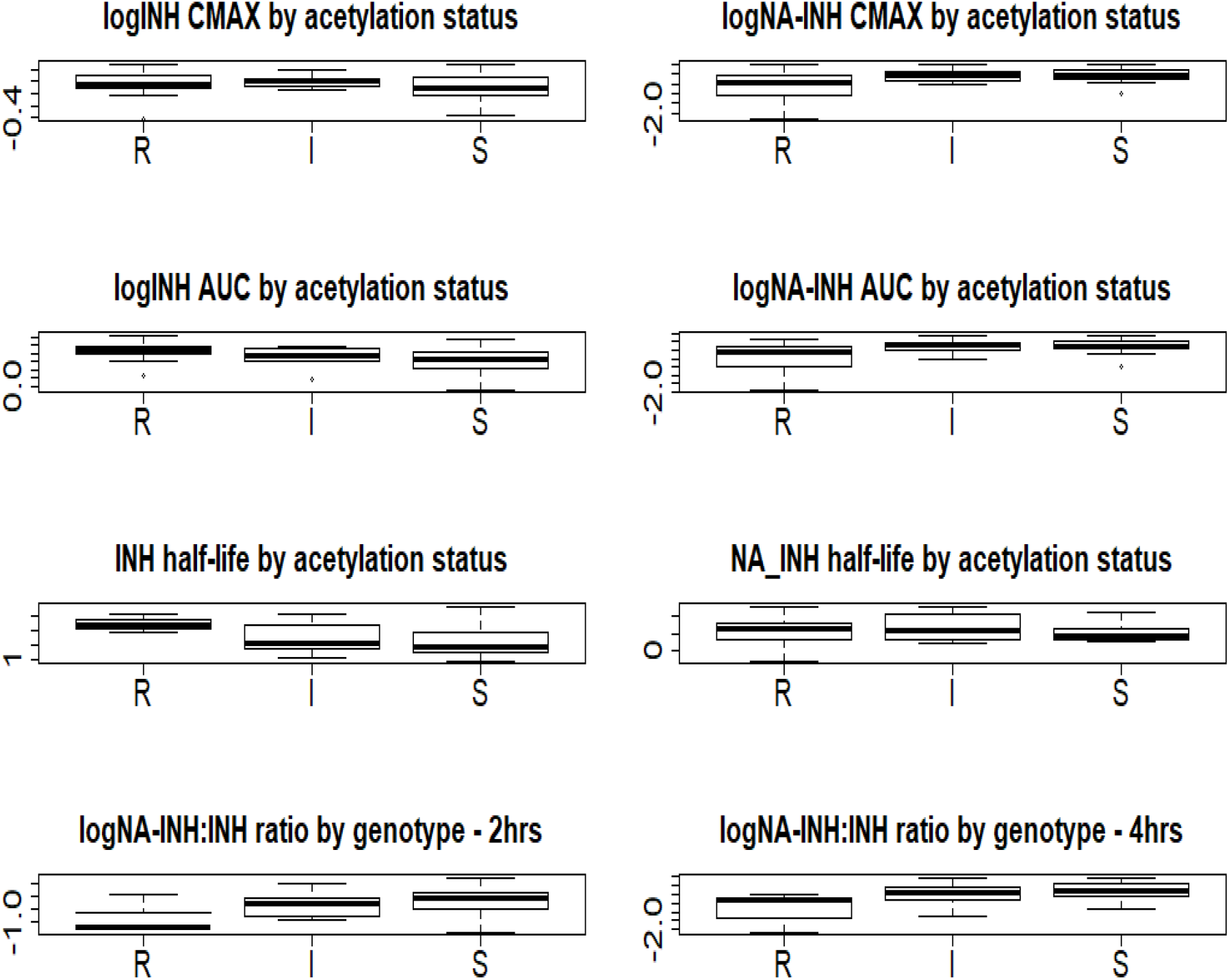
Box plots for study 1: representing median (solid line), interquartile range (box) and range (whiskers) for the pharmacokinetic parameters; log_10_ maximum concentration (C_max_), log_10_ area under the time-concentration curve (AUC_0-∞_), of isoniazid (INH) and *N*-acetyl-INH (AcINH) stratified by acetylator status and logAcINH to logINH ratio at 2 and 4 hours stratified by acetylator genotype.

**Figure 3:**
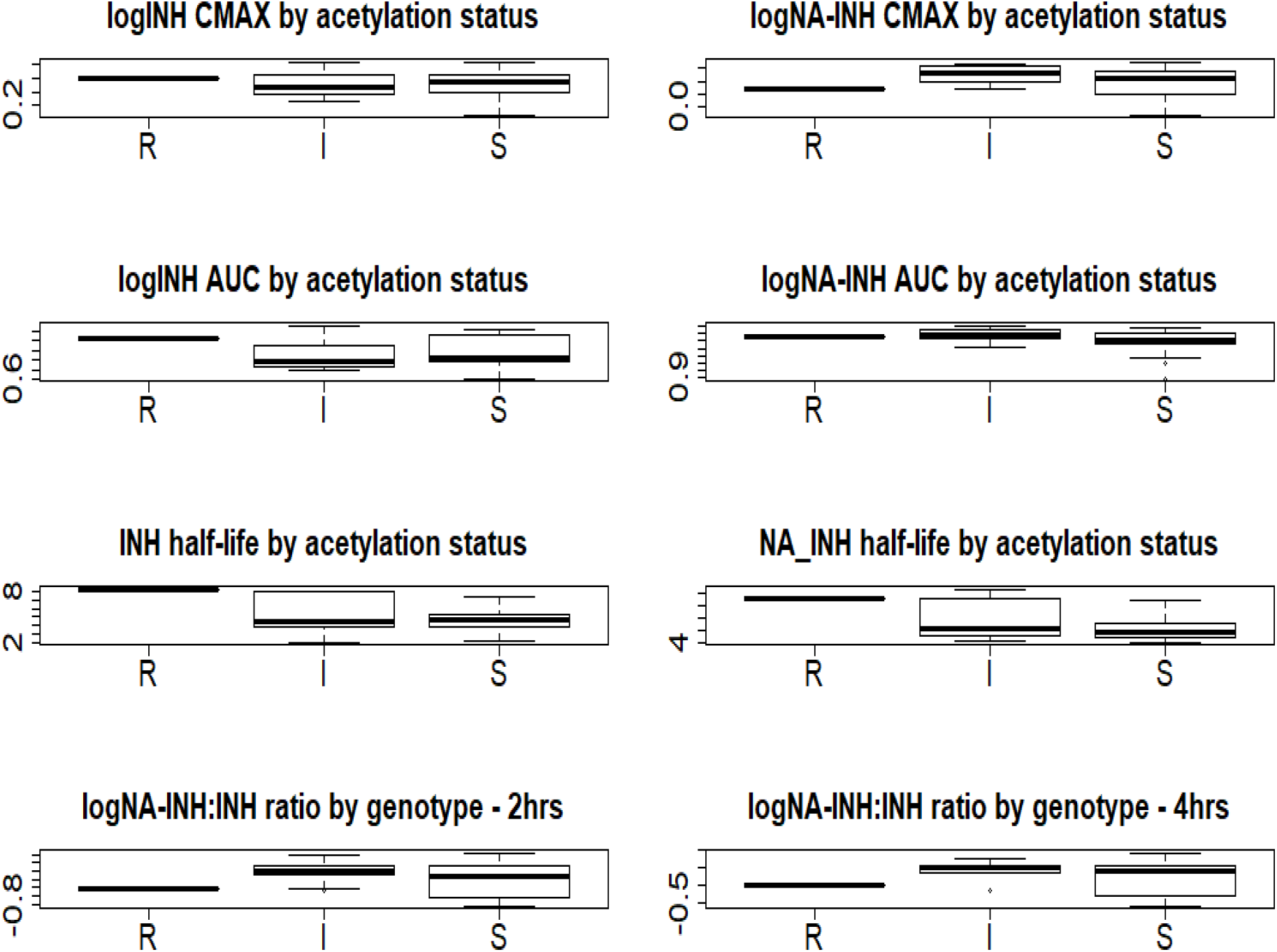
Box plots for study 2: representing median (solid line), interquartile range (box) and range (whiskers) for the pharmacokinetic parameters; log_10_ maximum concentration (C_max_), log_10_ area under the time-concentration curve (AUC_0-∞_), of isoniazid (INH) and *N*-acetyl-INH (AcINH) stratified by acetylator status and logAcINH to logINH ratio at 2 and 4 hours stratified by acetylator genotype.

### Hepatic adverse events

There were no grade 3 and 4 hepatic adverse events in Study 1 and only 1 grade 4 hepatic event was reported from the only participant with rapid genotype in Study 2. Although there were more grade 1 hepatic adverse events among the slow genotype participants, as shown in table 9, the difference was not statistically significant between genotypes; p=0.203 in Study 1, and 0.276 in Study 2.

**Table 9:** Participants with any hepatic adverse events. Hepatic adverse events from the two studies include a combination of elevated Aspartate, Aminotransferase (AST), Alanine Aminotransferase (ALT), Alkaline Phosphatase (ALP), Gamma-Glutamyl Transferase and total bilirubin.

|  | Study 1 |  |  |  | Study 2 |  |  |  |
| --- | --- | --- | --- | --- | --- | --- | --- | --- |
| AE Grade | Rapid N(%) | Intermediate N(%) | Slow N(%) | Total N(%) | Rapid N(%) | Intermediate N(%) | Slow N(%) | Total N(%) |
| Grade 1 | 7 | 9 | 25 | 41 | 0 | 5 | 10 | 15 |
| Grade 2 | 2 | 0 | 0 | 2 | 0 | 1 | 1 | 2 |
| Grade 3 | 0 | 0 | 0 | 0 | 0 | 2 | 1 | 3 |
| Grade 4 | 0 | 0 | 0 | 0 | 1 | 0 | 0 | 1 |
| Total | 9 (20.9) | 9 (20.9) | 25 (61) | 43 (100) | 1(4.8) | 8(30.1) | 12(57.1) | 21(100) |

## Discussion

We investigated the *NAT2* genotype, isoniazid and AcINH PK of black Zulu South Africans living with HIV from Durban and surrounding areas. We found that most individuals were of SA (52.5%) or IA (35.8%) genotype, with only a small number of RA genotype (11.7%). The proportions of the deduced acetylator phenotypes in our population was broadly similar to other African and Caucasian populations (36, 43, 57, 58) but differed from those previously reported from within other black ethnic groups within Southern Africa. For example, Werely *et al* found that IA genotypes dominated in the Xhosa cohort, with SAs only 30% (45). Our results were comparable to a recent Study by Naidoo et al. in patients from the same geographic area reported 34% SA, 43% IA and 18% RA (59).

There was a high prevalence of the *NAT2*5* allelic group in our population, accounting for the slow acetylator genotype. In well studied Caucasian and Asian populations, four variants; *NAT2*4* (wild type, rapid), *NAT2*5B, NAT2*6A*, and *NAT2*7B* (all slow), account for most *NAT2* alleles. In Asian populations there are generally a higher proportion of wild type *NAT2*4* alleles and few *NAT2*5B* alleles, and this difference largely accounts for the much lower prevalence of RAs in non-Asian populations. Consistent with other studies in Sub-Saharan African populations, the wild-type *NAT*4* allele was far less prevalent and variant alleles were far more diverse in our Study. In our population, the *NAT2*5B* allele was relatively rare in comparison to two studies in the black population from Western Cape and North West Province. (45, 60). However, in contrast to these populations, there were a diversity of other *NAT2*5* alleles, including a much higher prevalence of the rare *NAT2*5J* allele (17.5%) and the poorly characterised *NAT2*5K* allele (10.4%). The *NAT2*6A* and *NAT2*7B* alleles, common in Caucasian and Asian populations, were not seen in our cohort. In Caucasian and Asian populations, rapid *NAT2*12* alleles are rarely seen, where as in populations in sub-Saharan Africa the *NAT2*12A* allele is reported at much higher frequencies(35). In our Study the *NAT2*12A* allele did indeed comprise 5.8% of alleles seen but we saw a much higher frequency of the *NAT2*12C* allele (21.2%), in contrast to other Southern African cohorts(10, 45, 60, 61).

Isoniazid C_max_ and AUC_0-∞_ demonstrated considerable variability between individuals in both studies and almost all participants in Study 1 and almost half of the participants in Study 2 had a C_max_ below the lower limit of the target range(56). Low isoniazid concentrations during TB treatment are concerning because it is postulated they may lead to poorer treatment outcomes, or the generation of isoniazid resistance, the likely first step in the evolution of multi-drug resistant TB (MDR TB). However, the evidence for either of these concerns is mixed and in this setting the prevalence of INH mono-resistance is relatively low.

There was a marked difference in PK measures between the two studies analysed, with Study 1 having much lower measures than Study 2. There are several reasons that could have contributed to this difference. The difference in isoniazid dosing could explain the lower PK measures, where Study 1 used the FDC dosing as per WHO recommended weight bands, leading to almost half the participants receiving doses <300 mg, as previously reported(49). All participants in Study 2 received 300 mg doses of isoniazid irrespective of weight. Although the samples of Study 1 did not appear to deteriorate during the 5 years between first analysis and subsequent analyses for this study, differences in processing and storage between the studies cannot be excluded. Figure 3 shows the INH and AcINH at different time points. Based on this, the phenotype of the study participants is generally more intermediate/rapid than what the predominant slow genotype suggests, which is in contrast to other studies reporting HIV patients having a tendency towards slow phenotype (62).

We identified no statistically significant difference by *NAT2* genotype in a variety of PK measures, hence in this cohort we found poor correlation between *NAT2* genotype and phenotypic acetylation of isoniazid. Previous studies in other populations have shown good correlation between *NAT2* genotype and isoniazid PK, suggesting that *NAT2* genotyping could be used as a parsimonious way to risk-stratify patients and personalise dosing of isoniazid in an attempt to maximise efficacy whilst minimising toxicity. There are significant practical difficulties to implementing these approaches in this setting, but our data suggest that in this population *NAT2* genotyping will not be helpful in guiding TB therapy. A lack of concordance between genotypic and phenotypic measures of INH acetylation has been reported previously in HIV positive cohorts (63) (64). It is likely that in this cohort, as in others, other non-genetic factors are more or equally important than *NAT2* genotype. Jones et al found that infection with HIV or stage of HIV infection may alter Phase I and II drug metabolising enzyme (DME) activity in their study on 17 HIV infected participants at different levels of immunosuppression (65). They found that HIV infection was related to an increase in variability of these DMEs. Whilst additional pathways, aside from *NAT2* genotype, have been implicated in hepatotoxicity of isoniazid-containing TB treatment regimens, it is not clear that these pathways alter isoniazid PK and thus could account for the lack of genotypic and phenotypic concordance in this study.

Although there were more hepatic adverse events among the SA, there was no statistical association between genotype and hepatotoxicity in the two studies, with only 1 patient who was a RA having a grade 4 hepatic adverse event and 2 others who were IA having grade 3 hepatic adverse events.

In our study, participants received pyridoxine and cotrimoxazole with the ATT in Study 2, but not in Study 1 as we used the samples collected on day 1 for this analysis when only ATT was given. As both INH and sulfamethoxazole are inhibitors of CYP2C9, this could be one of the reasons for the variations noted. INH also inhibits CYP3A4, which is induced by rifampicin, this interaction has not proven significant except when it relates to hepatotoxicity (66, 67). That the combination of INH and rifampicin leads to an increased risk of hepatotoxicity, has been reported in other studies. In our Study 2, isoniazid was given with Rifabutin which is a less potent hepatic enzyme inducer, which therefore should have less interaction with INH (68). Considering the limited effect on hepatotoxicity, the effect of CYP2E1 was not evident in our study. We cannot confirm or exclude the effect of these CYP450 enzymes on INH metabolism in these participants.

In our study samples were stored at −80^0^ Celsius and loss of compound due to storage would have been minimal (69), although studies have not reported on plasma samples stored longer than 5 weeks, nor sample integrity for the metabolite, AcINH.

## Conclusion

Amongst black Zulu TB-HIV coinfected South African patients, most had slow or intermediate *NAT2* genotype. There was a diversity of specific *NAT2* alleles of a pattern differing from previously studied cohorts in other settings. Despite the rarity of rapid acetylator genotypes, INH PK was variable and a substantial proportion of individuals failed to attain minimum efficacy targets. Importantly *NAT2* genotype did not explain PK variability in this cohort or the low C_max_, which suggests that other factors could be influencing isoniazid bioavailability and metabolism, which require further elucidation.

## Acknowledgements

The study was sponsored by the Special Programme for Research and Training in Tropical Diseases, World Health Organization and United States Agency for International Development (USAID, Umbrella grant no. AAG-G-00-99-00005). The European and Developing Countries Clinical Trials Partnership (EDCTP) supplied supplementary funding of the PhD. We also acknowledge the generous donations of antiretroviral drugs from two major pharmaceutical companies, GlaxoSmithKline (UK) and Merck (USA), without which the study would not have been conducted. Study 2 was funded by Agence Nationale de Recherche Sur le Sida et les Hépatites Virales. Wellcome Trust (203919/Z/16/Z) supported one of the authors. These sponsors had nothing to do with study conduct. We would also like to thank the study staff for their hard work and patients for their involvement in this study.

